# A complete mitochondrial genome-based phylogeny of gooseneck barnacles confirms the sister group relationship between the eastern *Pacific Pollicipes* elegans and the eastern Atlantic *P. pollicipes*

**DOI:** 10.1101/2025.02.18.638925

**Authors:** Ruben Alfaro, Guillermo Mantilla, Alan Marín

**Affiliations:** Ecobiotech Lab S.A.C. Trujillo, Perú; Laboratorio de Genética, Fisiología y Reproducción, Facultad de Ciencias, Universidad Nacional del Santa, Chimbote, Perú

**Keywords:** Mitogenome, Phylogenetics, Cirripedes, Congener, Short Read Sequences

## Abstract

**Background:** Gooseneck barnacles are a fascinating invertebrate group and their taxonomic classification has attracted the attention of several specialists, including Charles Darwin, who was the first to note the close relationship between two species of gooseneck barnacles found on the eastern coasts of the Pacific, *Pollicipes elegans*, and the Atlantic, *P. pollicipes*. Additionally, a third species from the eastern Pacific, *P. polymerus*, was believed to have diverged earlier from the group.

**Methods and results:** Here, we used genomic short-read sequences publicly available in GenBank to assemble the first complete mitochondrial genome of *P. elegans*. We conducted a phylogenetic analysis to assess the relationships among its related species and other cirripede species from the superorder Thoracicalcarea. Our phylogenetic findings based on complete mitogenomes reinforce the close relationship between *P. elegans* and *P. pollicipes*, with *P. polymerus* emerging as a basal branch, indicating an earlier divergence from the group.

**Conclusions:** Our analysis provided the first robust phylogenetic assessment of *P. elegans*, strongly supporting its close relationship with *P. pollicipes*. Further efforts should aim to sequence the complete mitogenomes of other species within the order Pollicipedomorpha. This endeavor will enable a more comprehensive phylogenetic analysis of this group, whose classification remains a subject of debate.

## Introduction

The genus *Pollicipes* (Pollicipedomorpha: Pollicipedidae), commonly known as gooseneck barnacles, includes sessile pedunculated cirripedes that live in marine intertidal and subtidal rocky shores. Gooseneck barnacles not only represent species of economic significance across their distribution ranges but also serve as important marine biosensors [1] and have the potential to be a source of new adhesive materials for bioengineering [2]. Currently, there are four extant species within this genus, which are found along the eastern coastlines of the Atlantic (*P. pollicipes* and *P. caboverdensis*) and the Pacific oceans (*P. polymerus* and *P. elegans*) [3-5].

The taxonomic classification of gooseneck barnacles has garnered the attention of several specialists, including Charles Darwin, who was the first to report that *P. elegans* and *P. pollicipes* are more closely related to each other than they are to *P. polymerus* [6]. Darwin’s findings have received support from recent morphological and molecular phylogenetic analyses using partial gene sequences [4, 7, 8]. However, those phylogenetic results lacked strong statistical support, highlighting the need for more robust phylogenetic tools, such as mitochondrial genome [9].

To date, complete mitogenomes are available only for *P. pollicipes* and *P. polymerus*. In contrast, *P. elegans* has received less attention and its mitochondrial genome has not yet been sequenced. In this context, this study aims to assemble the complete mitogenome sequence of *P. elegans* using publicly available short read sequences (SRA) from the GenBank database. Additionally, we use the nucleotide sequence information of all protein-coding genes of the mitogenome of *P. elegans* to conduct a comprehensive phylogenetic analysis alongside the mitogenomes of other cirripede species available in GenBank.

## Materials and methods

### Genomic NGS reads mining, mitogenome assembly, and gene annotation

The NGS reads used to assemble the mitochondrial genome of *P. elegans* were retrieved from GenBank SRA repository (SRA accession SRR955082, Submitter: Clemson University). The *P. elegans* specimen (BioSample SAMN02324236) was collected from Punto Gaspareno, near Cabo San Lucas, Baja California, Mexico (coordinates 23°10’58.09”N, 110° 8’26.51”W) as part of the BioProject PRJNA216107. The raw reads were subjected to quality control using FastQC and trimmed with BBDuk as implemented in Geneious Prime V. 2025.0 (Biomatters Ltd., Auckland, New Zealand).

We conducted two different approaches namely “map to reference” and a hybrid “map to reference/de novo” approach. In the first approach, trimmed reads from *P. elegans* were assembled against the complete mitogenome reference sequences of *P. pollicipes* (GenBank accession NC_073544; 15,148 bp), using the “Map to Reference” tool in Geneious Prime with default parameter settings recommended by the software developers. For the second approach, the contig obtained from the first assembly approach were dissolved and reassembled “de novo” following the parameter settings recommended by the software developers selecting the “circularize contigs” option. The consensus sequences were extracted and verified manually using the Geneious Prime alignment tool.

Gene annotation was conducted using the “Annotate and Predict” tool of Geneious Prime by comparing the mitogenome of *P. elegans* to other complete mitogenomes of closely related species. The positions of the start and stop codons from the protein-coding genes (PCGs) were verified using the Open Reading Frame Finder tool (available at https://www.ncbi.nml.nih.gov/orffinder/) with the invertebrate mitochondrial genetic code. The locations of the transfer RNA genes (tRNAs) were verified using tRNAscan-SE [10]. The circular mitogenome were generated with Organellar Genome Draw “OGDRAW” version 1.3.1 [11] and edited with Inkscape macOS version 1.4 [12].

### Bayesian phylogenetic analysis

The Bayesian phylogenetic analysis was conducted using the 13 PCGs of the mitogenome assembled herein along with those of species of the superorder Thoracicalcarea available in GenBank, comprising 16 species from 9 families within 4 orders. Nucleotide sequences from each PCG were multialigned separately using MEGA 7 [13] and concatenated with SeaView version 5.0.4 [14], totaling a final matrix of 10,971 nucleotides. Models of nucleotide evolution for each one PCG were obtained using jModelTest 2 [15] under the Bayesian Information Criterion (BICc). Levels of substitution saturation at single codon positions from each PCG were assessed with Xia’s method implemented in DAMBE7 [16]. The ATP8 gene displayed saturation at the first and third codon positions, while the ATP6, CO2, CO3, CYTB, ND2, ND3, ND4L, ND5, and ND6 genes showed clear signs of saturations at their third codon positions and thereby the Bayesian analysis was performed with the data partitioned by gene and by codon position excluding the aforementioned codon positions. The Bayesian phylogenetic was conducting using MrBayes 3.2.7 [17] as implemented on the CIPRES science gateway 3.3 server [18], using two independent runs of four Markov chains each, conducted for 10 million generations, with samplings frequency of 1000. The first 25% of the sampled trees were discarded as burn-in. The final consensus phylogenetic tree was visualized using Figtree version 1.4.4 (http://tree.bio.ed.ac.uk/software/figtree/).

## Results and discussion

In this study, we successfully assembled the first complete mitogenome sequence of *P. elegans*. The nucleotide sequence data reported is available in the Third Party Annotation Section of the DDBJ/ENA/GenBank databases under the accession number TPA: BK070039. The mitogenome has a total length of 15,133 bp and includes the typical 13 PCGs, two rRNAs, and 22 tRNAs (Table 1 and Fig. 1). The majority of the PCGs (6 genes, 46.2%) utilizes the start codon ATG, while three genes (23.1% each) use the start codons ATT or GTG, and only 1 gene (7.6%) employs the ATA codon. Most of the PCGs (11 genes, 84.6%) terminate with the stop codon TAA, whereas the alternative stop codon TAG was found in only 2 genes. Incomplete stop codons (“T--” or “TA-“), which are completed posttranscriptionally by the addition of 3’ A residues by polyadenylation of the mRNAs [19], were identified in 6 of the PCGs (Table 1).

**Table 1.** Complete mitochondrial annotation and number of amino acids for each gene identified in the mitochondrial genome of *Pollicipes elegans*. The plus “+” and minus “-” symbols represent the heavy and light strands respectively. NCR: non-coding region.

| Gene | Strand | Location |  | Size (bp) | Anticodon | START codon | STOP codon | Amino acids |
| --- | --- | --- | --- | --- | --- | --- | --- | --- |
|  |  | Start | End |  |  |  |  |  |
| Lys (K) | - | 1 | 64 | 64 | UUU | — | — | — |
| Gln (Q) | - | 86 | 153 | 68 | UUG | — | — | — |
| Ile (I) | + | 268 | 336 | 69 | GAT | — | — | — |
| Met (M) | + | 336 | 399 | 64 | CAT | — | — | — |
| ND2 | + | 400 | 1,397 | 998 | — | ATG | TA- | 132 |
| Trp (W) | + | 1,398 | 1,461 | 64 | UCA | — | — | — |
| CO1 | + | 1,456 | 3,003 | 1,548 | — | ATA | TAA | 515 |
| Leu (L) | + | 3,008 | 3,075 | 68 | UAA | — | — | — |
| CO2 | + | 3,078 | 3,759 | 682 | — | ATG | T-- | 227 |
| Asp (D) | + | 3,760 | 3,825 | 66 | GUC | — | — | — |
| ATP8 | + | 3,826 | 3,984 | 159 | — | ATT | TAA | 52 |
| ATP6 | + | 3,978 | 4,642 | 665 | — | ATG | TA- | 221 |
| CO3 | + | 4,643 | 5,429 | 787 | — | ATG | T-- | 262 |
| Gly (G) | + | 5,430 | 5,491 | 62 | UCC | — | — | — |
| ND3 | + | 5,492 | 5,843 | 352 | — | ATT | T-- | 117 |
| Arg (R) | + | 5,844 | 5,906 | 63 | UCG | — | — | — |
| Asn (N) | + | 5,909 | 5,972 | 64 | GUU | — | — | — |
| Ala (A) | + | 5,973 | 6,033 | 61 | UGC | — | — | — |
| Glu (E) | + | 6,034 | 6,097 | 64 | UUC | — | — | — |
| Ser (S) | + | 6,098 | 6,153 | 56 | GCU | — | — | — |
| Phe (F) | - | 6,170 | 6,231 | 62 | GAA | — | — | — |
| ND5 | - | 6,227 | 7,933 | 1,707 | — | GTG | TAA | 568 |
| His (H) | - | 7,934 | 7,999 | 66 | GUG | — | — | — |
| ND4 | - | 8,000 | 9,329 | 1,330 | — | ATG | T-- | 443 |
| ND4L | - | 9,323 | 9,616 | 294 | — | GTG | TAA | 97 |
| Pro (P) | - | 9,617 | 9,680 | 64 | UGG | — | — | — |
| Thr (T) | + | 9,689 | 9,752 | 64 | UGU | — | — | — |
| ND6 | + | 9,753 | 10,238 | 486 | — | ATT | TAA | 161 |
| CYTB | + | 10,238 | 11,377 | 1,140 | — | ATG | TAG | 379 |
| Ser (S) | + | 11,376 | 11,445 | 70 | UGA | — | — | — |
| Tyr (Y) | - | 11,460 | 11,522 | 63 | GUA | — | — | — |
| Cys (C) | - | 11,532 | 11,595 | 64 | GCA | — | — | — |
| ND1 | - | 11,594 | 12,520 | 927 | — | GTG | TAG | 308 |
| Leu (L) | - | 12,521 | 12,586 | 66 | UAG | — | — | — |
| 16S rRNA | - | 12,677 | 13,994 | 1,318 | — | — | — | — |
| Val (V) | - | 13,997 | 14,061 | 65 | UAC | — | — | — |
| 12S rRNA | - | 14,062 | 14,833 | 772 | — | — | — | — |
| NCR | + | 14,834 | 15,133 | 300 | — | — | — | — |

**Fig. 1.**
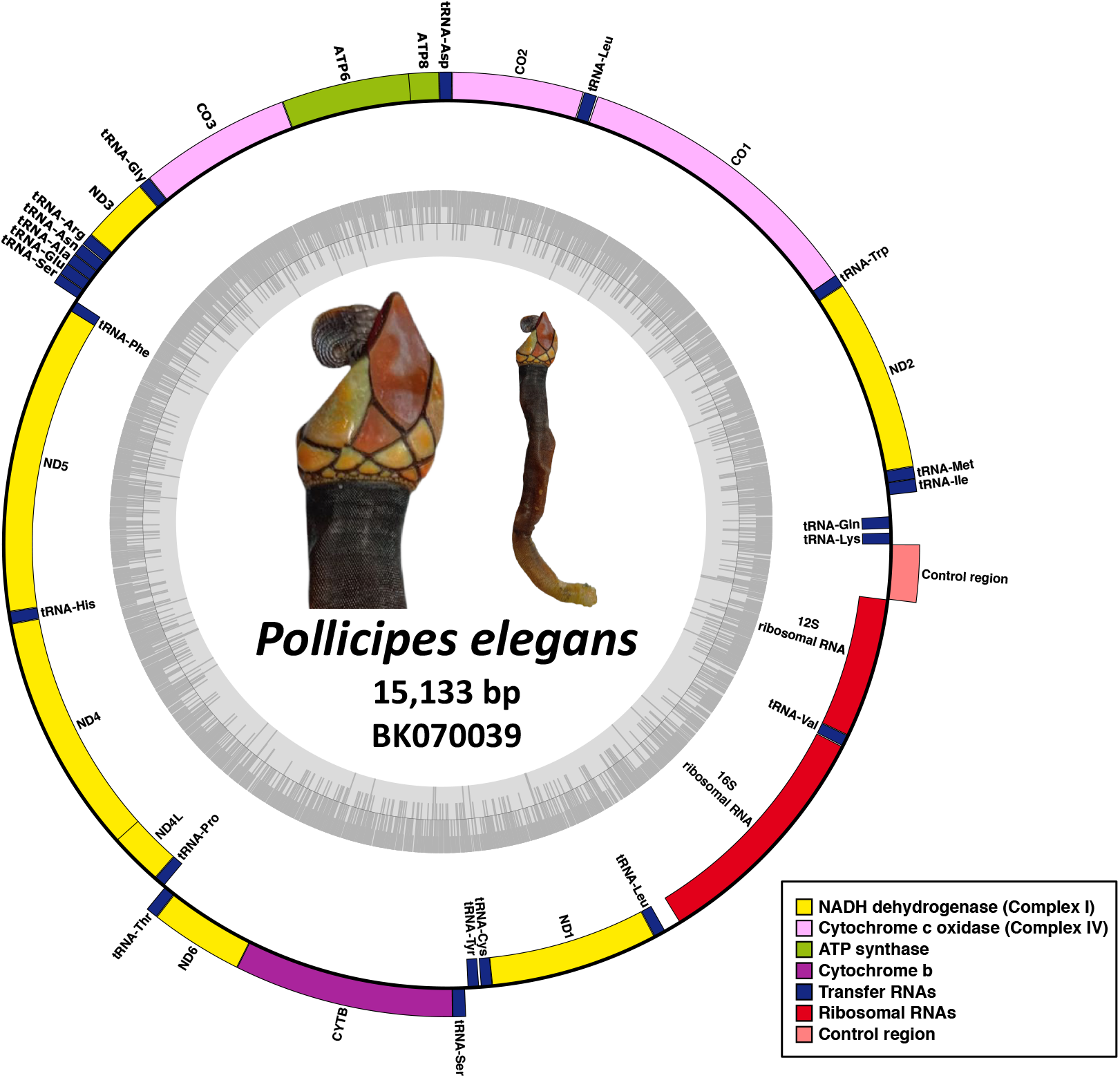
Circular mitochondrial genome organization of *Pollicipes elegans*. Genes located in inner circle (negative strand) are transcribed in a clockwise direction, while those on the outer circle are transcribed counterclockwise. The innermost circle of the GC content graph indicates a 50% threshold, with darker and lighter grey shades representing GC and AT content, respectively. The different gene types are illustrated with colored bars: ND dehydrogenase (yellow), cytochrome c oxidase (pink), ATP synthase (green), cytochrome b (purple), transfer RNA (blue), and ribosomal RNA (red). The control region is represented by a coral red bar.

A total of 22 genes are encoded in the heavy strand, which includes 9 PCGs and 13 tRNAs. In contrast, the light strand contains 15 genes, comprising 4 PCGs, the 2 rRNAs, and 9 tRNAs. Overall, the gene rearrangement of the mitogenome of *P. elegans* is identical to that of its closest relative *P. pollicipes* (GenBank accession NC_073544).

The results of the Bayesian phylogenetic analysis (Fig. 2) revealed a well-supported *Pollicipes* clade with a Bayesian posterior probability (BPP) = 1, confirming the close relationship between the eastern Pacific *P. elegans* and the eastern Atlantic *P. pollicipes*, which were grouped in a single subclade. In contrast, the other eastern Pacific species, *P. polymerus*, was placed in a separate ancestral branch within the *Pollicipes* group. These findings align with previous morphological and molecular studies, which concluded that *P. elegans* and *P. pollicipes* are the most recently diverged species of the genus *Pollicipes*, having separated from the Tethys Sea approximately 25 to 34 million years ago (mya) [4, 7]. Meanwhile, *P. polymerus* is believed to have a more ancestral origin from the northwestern margin of the Tethys Sea (now the northeastern Pacific), dating back about 55 to 65 mya [4, 7].

**Fig. 2.**
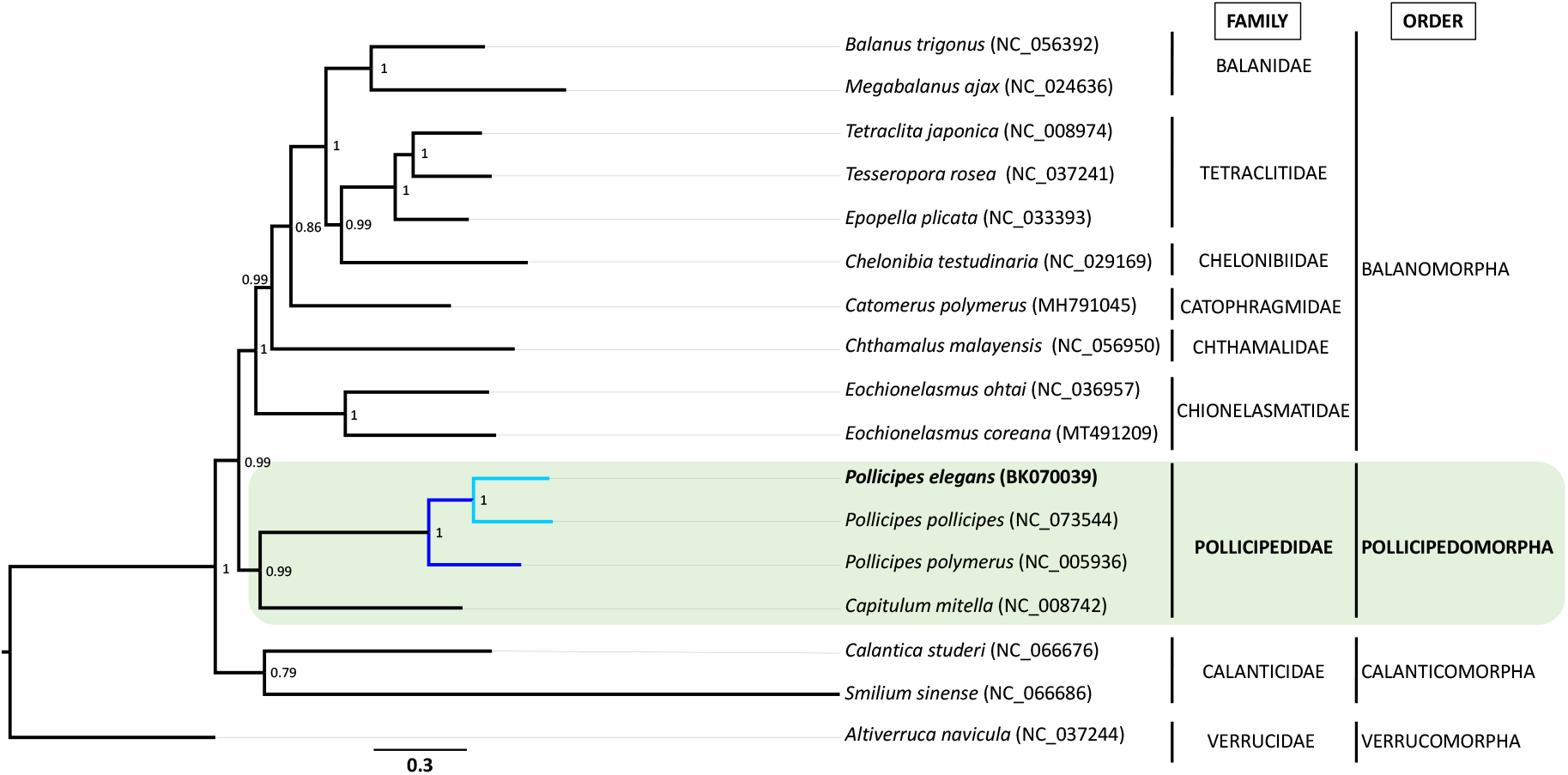
Bayesian phylogenetic tree constructed based on the concatenation of 13 mitochondrial protein-coding genes from 17 cirripede species (GenBank accessions are provided in parenthesis), including the newly determined mitogenome of *Pollicipes elegans* (family Pollicipedidae), which is highlighted in bold. The tree displays the subclades formed by *P. elegans* and *P. pollicipes* in sky blue color, while the branch containing to *P. polymerus* is shown in blue. Posterior probabilities are shown at the corresponding nodes. The deep-sea barnacle, *Altiverruca navicula*, was used as outgroup.

The Japanese goose barnacle, *Capitulum mitella*, which belongs to the family Pollicipedidae, was recovered at the basal position of the Pollicipedidae group (Fig. 2) with strong statistical support (BPP = 0.99). Previous studies have stablished a sister group relationship between *Pollicipes* and *Capitulum* based on morphological evidence and partial gene sequences [4, 20], as well as complete mitogenome analysis [21, 22]. However, a prior study that used a con- catenated dataset from partial nuclear (18S rDNA and Histone H3) and mitochondrial (CO1 and 12S rRNA) genes placed *C. mitella* outside the Pollicipedomorpha node [23]. These contrasting results emphasize the effectiveness and reliability of complete mitogenome phylogenetic analysis over partial gene sequences in resolving phylogenetic relationships. Our phylogenetic analysis established the order Pollicipedomorpha as a sister group to the order Balanomorpha, also with strong statistical support (BPP = 0.99), while Calanticomorpha was placed in a basal position (BPP = 1). This finding is consistent with previous mitogenome phylogenies reported in other studies [e.g. 22], suggesting that these three orders may share a common ancestor, with Calanticomorpha branching off first, followed by Pollicipedomorpha and Balanomorpha.

## Conclusions

Our analysis provided the first robust phylogenetic assessment of *P. elegans*, strongly supporting the close relationship between *P. elegans* and *P. pollicipes*, a connection that have been previously identified only through morphological characteristics and partial gene sequences. Further efforts are needed to sequence the complete mitogenomes of other species within the order Pollicipedomorpha, such as *Anelasma squalicola* and *P. caboverdensis* (Pollicipedidae) as well as *Lithotrya* spp. (Lithotryidae). Obtaining these mitogenome sequences will facilitate a comprehensive phylogenetic analysis of this intriguing group of crustaceans. Additionally, population genetic studies and assessments of genetic diversity are needed for *P. elegans* from northern Peru, as these populations have been subjected to overexploitation due to their economic value [9].

## Notes

### Competing Interest Statement

The authors have declared no competing interest.

